# A Rat Model of Somatosensory-Evoked Reflex Seizures Induced by Peripheral Stimulation

**DOI:** 10.1101/506352

**Authors:** Aleksandra Bortel, Ze Shan Yao, Amir Shmuel

## Abstract

**Objective:** We introduce a novel animal model of somatosensory stimulation-induced reflex seizures which generates focal seizures without causing damage to the brain.

**Methods:** Specifically, we electrically stimulated digits or forepaws of adult rats sedated with dexmedetomidine while imaging cerebral blood volume and recording neurophysiological activity in cortical area S1FL. For the recordings, we either inserted a linear probe into the D3 digit representation or we performed surface electrocorticography (ECoG) recordings.

**Results:** Peripheral stimulation of a digit or the forepaw elicited seizures that were followed by a refractory period with decreased neuronal activity, or another seizure or a normal response. LFP amplitudes in response to electrical pulses during the seizures (0.28 ± 0.03 mV) were higher than during normal evoked responses (0.25 ± 0.05 mV) and refractory periods (0.2 ± 0.08 mV). Seizures generated during the stimulation period showed prolonged after-discharges that were sustained for 20.9±1.9 s following the cessation of the stimulus. High-frequency oscillations were observed prior to and during the seizures, with amplitudes higher than those associated with normal evoked responses. The seizures were initially focal. Optical imaging of the cerebral blood volume response showed that they propagated from the onset zone to adjacent cortical areas, beyond the S1FL representation of the stimulated digit or forepaw. The spatial extent during seizures was on average 1.74 times larger during the stimulation and 4.1 times following its cessation relative to normal evoked responses. Seizures were recorded not only by probes inserted into cortex but also with ECoG arrays (24.1±5.8 seizures per rat) placed over the dura matter, indicating that the seizures were not induced by damage caused by inserting the probes to cortex. Stimulation of the forepaw elicited more seizures (18.8±8.5 seizures per rat) than stimulation of a digit (1.7±0.7). Unlike rats sedated with dexmedetomidine, rats anesthetized with urethane showed no seizures, indicating that the seizures may depend on the use of the mild sedative dexmedetomidine.

**Significance:** Our proposed animal model generates seizures induced by electrical sensory stimulation free of artifacts and brain damage. It can be used for studying the mechanisms underlying the generation and propagation of reflex seizures and for evaluating antiepileptic drugs.

**HIGHLIGHTS:**

- Peripheral stimulation of the rat forepaw or digits induces seizures
- Seizures are evoked with no direct application of convulsants, electro-stimulation or damage to the brain
- Seizures are focal at onset, then spread beyond the spatial representation of the digit or forepaw
- Seizures persist following the cessation of the stimulus
- Proposed animal model may support the study of reflex seizures and improving therapeutic interventions

## 1. INTRODUCTION

Reflex seizures are seizures elicited by specific sensory stimulation. These seizures may be triggered by various types of stimuli that are patient-specific (Panayiotopoulos, 2005; Sala-Padro et al., 2015; Striano et al., 2012; Wolf, 2015). In humans, reflex seizures can be induced by simple somatosensory stimulation, such as a cutaneous sensorial stimulus (Koepp et al., 2016; Loscher, 1999; Sarkisian, 2001; Wolf, 2015). Touch-based somatosensory-evoked seizures, termed Touch Induced Seizures (TIS), are associated with cortical hyper-excitability and are classified as reflex seizures induced by a simple stimulus (Sala-Padro et al., 2015; Striano et al., 2012; Wolf, 2015). In humans, TIS can be evoked by specific movements such as skin friction, touching, tapping, tooth brushing, or stimulation of the external ear canal (Italiano et al., 2014; Striano et al., 2012). Electrical stimulation of peripheral nerves can also induce somatosensory-evoked seizures (Deonna, 1998). Usually, TIS are focal seizures with no impairment of consciousness (Deonna, 1998; Sala-Padro et al., 2015; Striano et al., 2012) and may have only electrographic display without any overt clinical manifestations (Panayiotopoulos, 2005). Here we introduce a novel animal model of somatosensory-induced reflex seizures which generates sensory stimulation-induced focal seizures without causing damage to the brain.

To date, only genetically modified epilepsy-prone species have been described as models for studying reflex seizures, specifically photosensitive and audiogenic reflex seizures (Italiano et al., 2016; Kandratavicius et al., 2014; Sarkisian, 2001). Current animal models that are seizure-susceptible in response to photic and/or audiogenic stimulation include the photosensitive baboon (Papio papio), the audiogenic epilepsy prone rat (GEPR) and DBA/2 mice (Kandratavicius et al., 2014; Sarkisian, 2001). However, to date, no animal model has been introduced for inducing seizures by peripheral somatosensory stimulation. Here we introduce an animal model of somatosensory-evoked reflex seizures induced by electrical stimulation, without direct administration of electrical currents or convulsants to the brain.

After-discharges are epileptiform discharges that can be triggered by chemoconvulsants or direct electrical stimulation of different brain structures (Lesser et al., 2008; Racine, 1975) and are morphologically similar to spontaneous seizures (Blume et al., 2004). It was previously shown that repetitive intracranial electrical stimulation elicits self-sustained after-discharges (SSADs) in the sensorimotor cortex of cats (Dietzel et al., 1982; Steriade and Yossif, 1974). The cortical SSADs can be induced by rhythmic stimulation of subcortical structures such as the thalamic somatosensory relay nucleus or by the stimulation of the dorsal hippocampus (Mares et al., 1984). Our study observes epileptiform discharges that are similar to those reported in previous SSAD studies; however, our study can be distinguished from previous SSAD studies because we induce seizures by peripheral sensory stimulation rather than direct intracranial stimulation.

Electrical stimulation or topical administration of chemical compounds including pentylenetetrazol, tetanus toxin, and penicillin have been used to elicit seizures in animal models for many years (Engel, 2004; Kandratavicius et al., 2014; Sarkisian, 2001). However, these techniques bear several disadvantages. First, an electrical stimulation technique for inducing acute seizures, termed *electroshock-induced seizures*, modifies activity in the larger part of the brain. Second, all electrical stimulation techniques used to induce seizures - including electrical kindling - cause electrical artifacts that interfere with studying the spatiotemporal dynamics of neuronal activity. Third, the systemic administration of kainate and pilocarpine also cause damage to the brain by eliciting inflammation, activation of glial cells, and disruption of the blood-brain barrier (Sperk, 1994; Turski et al., 1989; van Vliet et al., 2016). Similarly, chemical injections into the brain lack spatiotemporal precision and cause neuronal damage and reactive gliosis (Osawa et al., 2013). This motivates the introduction herein of an artifact-free model of seizures in adult rats evoked by peripheral somatosensory stimulation of a digit or forepaw.

Seizure symptoms of epilepsy are often associated with highly modified cortical activity. Indeed, seizures induced by direct application of chemoconvulsants to rat brain slices lead to permanent changes in the somatosensory representation of the cortex (Borbely et al., 2006). Similarly, *in vivo* direct intracranial electrical stimulation of the rat corpus callosum and neocortex causes seizures and substantially modifies activity in the sensorimotor cortex (Flynn et al., 2010; Ozen et al., 2008; Vuong et al., 2011).

Changes in LFP frequency bands are informative for seizure description and prediction (Modur et al., 2011). For instance, in epileptic patients and animals, the power of pathological high frequency oscillations (HFOs) increase before seizure initiation (Jacobs et al., 2009; Modur et al., 2011; Staba, 2012) and reflect the activity of dysfunctional neural networks that sustain the epileptogenesis (Behr et al., 2014). Therefore, HFOs may serve as biomarkers of epileptogenic neural networks and predictors of seizure occurrence (Jefferys et al., 2012).

Here we demonstrate the induction of seizures in adult rats by peripheral electrical somatosensory stimulation of the digits or forepaw. To this end, we performed simultaneous intrinsic optical imaging and neurophysiological recordings. We characterize the seizures and the associated HFOs, and we analyze the conditions required to elicit them. We present the seizures’ spatial extent relative to normal sensory responses to peripheral electrical stimulation in cortical area S1FL. Our study introduces an artifact-free animal model of somatosensory-induced reflex seizures.

## 2. MATERIALS AND METHODS

### 2.1. Animals and surgical procedures

All procedures were approved by the animal care committees of the Montreal Neurological Institute and McGill University, and were carried out in accordance with the guidelines of the Canadian Council on Animal Care. Experiments were performed on 28 adult, 100-107 days old male Sprague-Dawley rats weighing 400-580 g. The rats were housed under controlled environmental conditions at 22 ±2° C with a 12h light/12h dark cycle (lights on from 7:00 a.m. to 7:00 p.m.) and received food and water ad libitum.

The rats were initially anesthetized with a solution of xylazine (10 mg/kg i.p.; Bayer Inc., Canada) and ketamine (50 mg/kg i.p.; Wyeth, Canada). They were then intubated and placed in a stereotaxic frame. The surgical procedure was performed under anesthesia with isoflurane (0.6-2%; Benson Medical Industries Inc., Canada). During the surgery, the rats were ventilated with a mixture of oxygen (50%) and medical air (50%). Physiological parameters, including temperature, heart rate, and PaO2 measured with pulse oximetry, were monitored continuously. The scalp was incised to expose the skull covering the primary somatosensory cortex (S1) of the left hemisphere. One stainless steel screw (2.4 mm in length) was fixed to the skull above the visual cortex of the right hemisphere and was used as a ground and reference. The area of skull overlying S1 was thinned until soft and transparent. We then performed an approximate 4 mm wide square-shaped craniotomy, centered on the forelimb representation in area S1 (S1FL) based on a stereotaxic atlas (AP 0.00 mm, ML ±4.50 mm, DV –2.5 mm) (Paxinos and Watson, 2005). The dura was resected, and a linear multi-contact probe was inserted into area S1FL. At the end of the surgical procedure, a silicon chamber was created around the craniotomy and filled with Hank’s Balanced Salt Solution (Invitrogen, Canada). Following the surgery, prior to starting the recordings, we switched to ventilating with a mixture of 20% oxygen and 80% medical air. At the same time, we injected a single dose of the morphine-related analgesic buprenorphine (0.04 mg/kg, s.c.; Schering-Plough, UK) and started administration of a sedative dexmedetomidine (0.075 mg/kg/h, s.c.; Pfizer Inc., Canada) that remained continuous throughout the recordings. To examine the effect of the anesthesia on the epileptic activity, we injected six animals with a single dose of an anesthetic urethane (1.2 g/kg, i.p.; Sigma-Aldrich, Canada) instead of administering dexmedetomidine and buprenorphine. The isoflurane administration was stopped following the administration of either dexmedetomidine and buprenorphine, or urethane.

### 2.2. Electrical stimulation of a single digit or forepaw

Electrical stimuli were delivered to the right digit D3, D5, or forepaw using a stimulator/isolator (A365, WPI, Sarasota, FL). For digit stimulation, we attached two electrodes (anode and cathode; Plastics One, Roanoke, VA) per digit, one on each opposing side of the digit. The digit stimulation electrodes were made of stainless steel and had impedance of approx. 14Ω. They had a rectangular shape 3.25 mm x 1.10 mm contacts. For forepaw stimulation, we inserted needle electrodes into the web spaces of digits 2/3 and 4/5 of the right forepaw.

We began our experiments by optical imaging of area S1FL to guide the insertion of the neurophysiology probe to the responding region. Stimulation runs for eliciting cerebral blood volume (CBV) based optical imaging responses (Figure 1) consisted of ten 12s-long trials, separated by inter-trial intervals of 12s during which no stimulus was delivered. Each stimulation trial began with 2s recordings of baseline activity, followed by 4s of stimulation and 6s of baseline activity. The 4s stimulus consisted of a train of electrical pulses delivered to D3, D5, or the forepaw at a frequency of 8 Hz. The stimulus intensity was 1 mA for forepaw and 1.5 mA for digit and the duration of each single electrical pulse was 1ms. The polarity of stimulation was switched in each pulse relative to the polarity in the preceding pulse.

**Figure 1.**
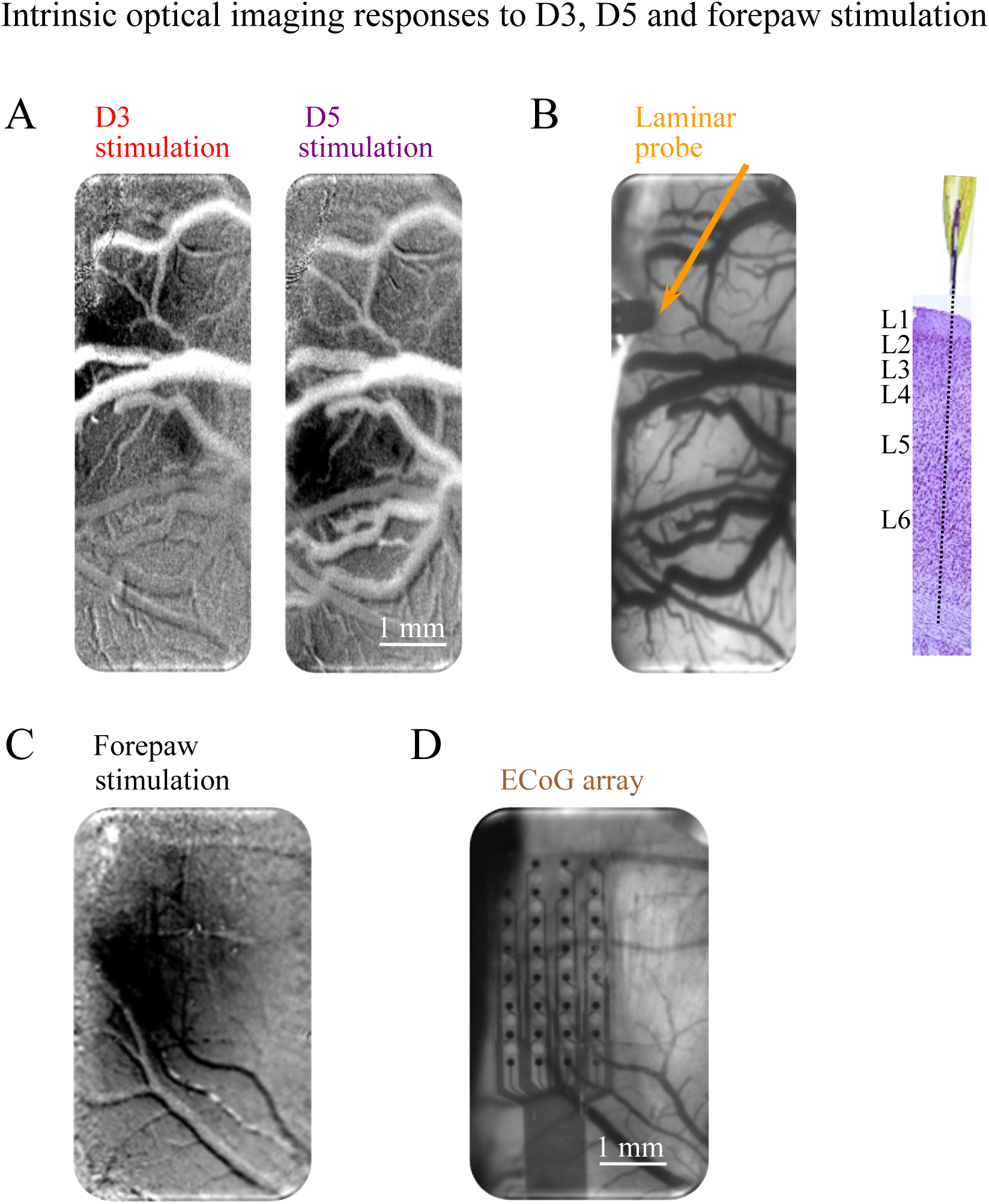
Optical imaging of intrinsic signal responses to D3, D5, and forepaw stimulation. **A.** The representations of D3 and D5 in area S1FL as identified by the blood volume response to separate stimulations of D3 and D5, imaged optically under the illumination wavelength of 530 nm. Dark regions showed increase in cerebral blood volume. **B.** The image to the left presents the cortical surface and the pial vessels, as imaged under the illumination wavelength of 530 nm. The orange arrow points to the site in which the probe (dark, seen to the left of the arrow) was inserted. A linear probe with 32 microelectrode contacts equally spaced at 100-µm intervals was inserted into the D3 representation of area S1FL. **C.** S1FL representation of the contralateral forepaw, obtained by the blood volume response to electrical stimulation, imaged optically under the illumination wavelength of 530 nm. **D.** An ECoG surface array with 32 contacts, positioned directly on top of the dura above area S1FL of the same brain imaged in C.

For inducing seizures, four runs of D3 stimulation, interleaved with four runs of D5 stimulation or eight forepaw stimulation runs were performed in each of the rats. These stimulation runs consisted of ten 35s-long stimulation trials, separated by ten 35s-long trials in which no stimulus was delivered (see time courses in Figures 2A and 3A). Each stimulation trial started with 5s recordings of baseline activity, followed by 10s of stimulation and 20s of baseline activity. The 10s stimulus consisted of a train of electrical pulses delivered to D3, D5, or the forepaw at a frequency of 8 Hz. The stimulus intensity was 2 mA and the duration of each single electrical pulse was 1ms. The polarity of stimulation was switched in each pulse relative to the polarity of the preceding pulse. Eight runs were performed in each of these rats.

**Figure 2.**
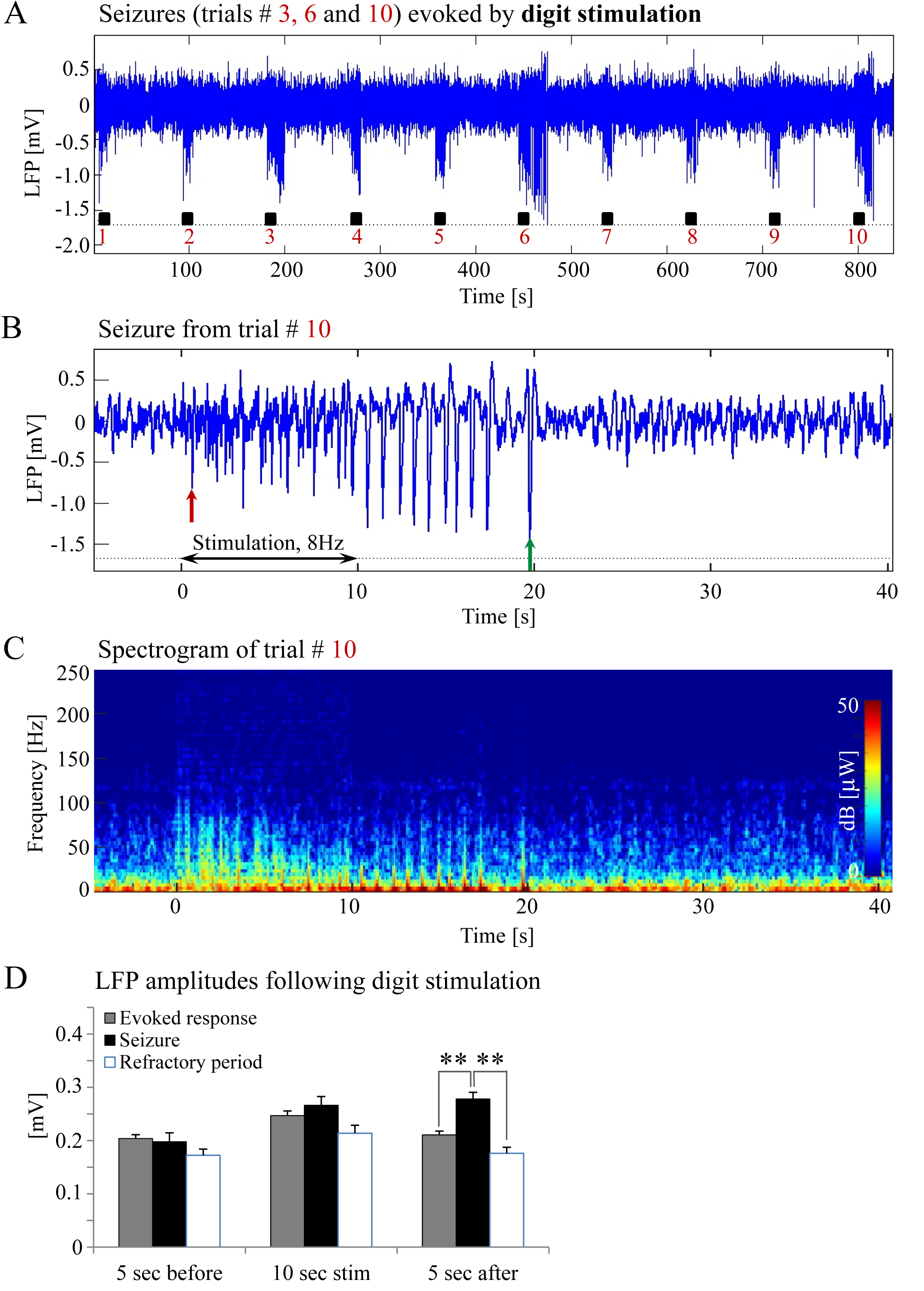
Evoked epileptic activity in response to electrical stimulation of D3. **A.** LFP recordings of ten trials, each with 10s-long stimulation. The stimulation periods are marked by black rectangles. Each 10s stimulus consisted of a train of electrical pulses delivered at 8 Hz to digit D3. Note that the 10s stimulation evoked seizures during the third, sixth, and tenth trials. **B.** The LFP (mean over all contacts spanning the cortical depth) demonstrates a seizure in area S1FL. The red and green arrows indicate the onset and termination, respectively, of a seizure. **C.** A spectrogram (power as a function of frequency and time) computed for the same seizure. **D.** The mean amplitudes of LFP recordings during baseline, stimulation and following stimulation for normal responses, seizures, and refractory periods. The mean amplitudes of LFP represents the mean of the absolute mean extra-cellular potentials, obtained by averaging over probe contacts (** p<0.001; Tamhane’s test).

**Figure 3.**
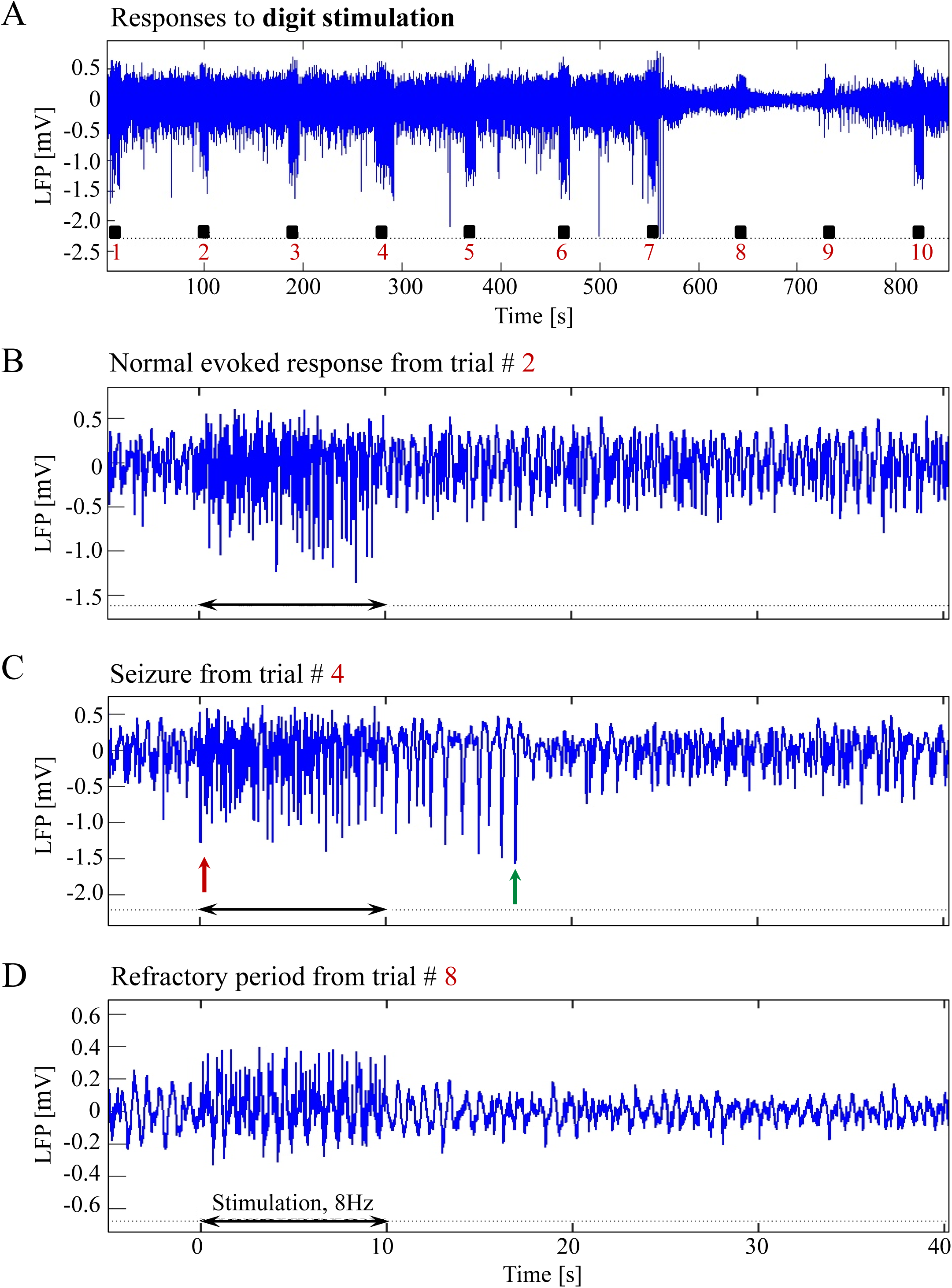
Responses to digit stimulation during normal response, seizure and refractory period. **A.** LFP responses obtained from area S1FL following D3 stimulation. The panel shows ten trials, each with 10s-long stimulation marked by a black rectangle. **B.** Normal evoked responses; the panel shows a magnification of the second trial from A. **C.** A typical activity pattern of an evoked seizure (magnification of the fourth trial from A). **D.** Refractory period (magnification of the eighth trial in A). The time bars for panels B, C and D are identical. Note that similar LFP patterns were observed following D3, D5, and forepaw stimulation.

### 2.3. Optical imaging of intrinsic signals

To target the linear probes and surface electrocorticography (ECoG) arrays to the region of S1 that responded to the electrical stimuli, we first imaged the cortical response using optical imaging of intrinsic signals (Grinvald et al., 1999). Cortical images were obtained using a 15-bit differential data acquisition system (Imager 3001, Optical Imaging Ltd., Rehovot, Israel) with a Pantera 1M60 camera (Teledyne Dalsa, Waterloo, Ontario, Canada) and a 60 mm lens. The camera was mounted above the optical chamber, with its optical axis perpendicular to the cortical surface. We focused the image on the surface of the gray matter and the cortical pial blood vessel in our region of interest (ROI). The ROI we imaged was centered on the atlas coordinates of area S1FL in the left hemisphere (AP 0.00 mm, ML ±4.50 mm, DV –2.5 mm; Paxinos and Watson (2005)), with a matrix of approximately 1000-by-500 pixels, 70–80 pixels/mm, and a 50 µm depth of field. The camera frame rate was set to 30 Hz. We imaged the CBV responses to D3, D5 (Figure 1A), and forepaw stimulation (Figure 1C) under illumination of a green LED light with a center wavelength of 530 nm (isosbestic point).

### 2.4. Electrophysiology

A linear probe with 32 contacts equally spaced at 100 µm intervals (Neuronexus technologies, Ann Arbor, MI) was inserted into the D3 representation within area S1FL using a manually driven micro-manipulator (David Kopf instruments, Tujunga, CA). All insertions, for both digit and forepaw stimulation studies, were made into the center of the D3 representation, which we estimated based on optical imaging of the CBV response to stimulation of D3 (see above). Before each insertion, we verified that the D3 representation we imaged was within area S1FL, as expected based on the rat brain atlas (Paxinos and Watson, 2005).

We monitored the probe insertion angle from multiple points of view, ensuring that the probe was approximately orthogonal to the local cortical surface. To assure coverage of the cortical depth by the contacts, the root mean square (RMS) of signals detected by the bottommost contact was monitored while driving the probe into the cortex. A decrease in RMS indicated when the bottommost recording contact had reached the cortical surface. The probe was then slowly lowered 2.7 mm deeper, spanning the entire thickness of the cortex. Approximately four contacts remained above cortex (within the silicon chamber filled with Hank’s Balanced Salt Solution), 21-22 contacts were within cortex and approximately six contacts were below cortex (in the white matter) (Figure 1B). The neurophysiological signals were sampled at 24,414.1 Hz using a multi-channel recording system (Tucker Davis Technologies, Alachua, FL).

In separate experiments, we tested whether the epileptic activity was possibly caused by damage to the cortex due to the insertion of the linear probe, which potentially could make the cortex prone to seizures. To this end, surface ECoG recordings were obtained from six rats. The surface ECoG array (NeuroNexus technologies, Ann-Arbor, MI) was centered on the forepaw representation in area S1FL, which was obtained by optical imaging of the CBV response to forepaw stimulation (Figure 1C). The array was placed on top of the dura, leaving the brain intact (Figure 1D). The neurophysiological signals were sampled at 24,414.1 Hz.

### 2.5. Experimental paradigms

Data were obtained from 28 rats under two anesthetic regimes (dexmedetomidine or urethane). Recordings of responses to either digit or forepaw stimulation were conducted using a linear probe or an ECoG array. Therefore, our data set is comprised of four experimental groups (Table 1):

**Table 1.** Average number of seizures, onset time, and duration of electrographic seizures.

| Group | Number of rats | Probe | Stimulation | Anesthesia regime | Mean # of Seizures | Onset [s] | Duration [s] |
| --- | --- | --- | --- | --- | --- | --- | --- |
| **1** | 10 | linear | digit | Dex / Bup | $1.70 \pm 0.70$ | $0.56 \pm 0.12$ | $30.63 \pm 2.56$ |
| **2** | 6 | linear | digit | Urethane | – | – | – |
| **3** | 6 | linear | forepaw | Dex / Bup | $18.83 \pm 8.49$ | $0.60 \pm 0.05$ | $31.50 \pm 1.83$ |
| **4** | 6 | ECoG | forepaw | Dex / Bup | $24.16 \pm 5.83$ | $0.88 \pm 0.09$ | $19.47 \pm 1.08$ * |

The first 5 columns present the 4 experimental groups in our study, using a linear probe or a surface ECoG array, digit or forepaw stimulation, and dexmedetomidine or urethane anesthesia. The sixth to eighth columns present the average number of seizures per animal, onset time relative to the stimulus onset, and duration of seizures. Results are expressed as a mean ± SEM. Note the statistically significant differences in seizure duration between probe and ECoG recordings (\* $p < 0.05$ ; Tamhane's test).

### 2.6. Preprocessing of electrophysiological data

The mean extracellular field potential was separated into the high-frequency portion (> 400 Hz), revealing multiunit activity (MUA), and the low-frequency portion (< 400 Hz), revealing the local field potential (LFP). Action potential detection and sorting based on the high-pass filtered signal were performed offline using ‘wave clus’ (Quiroga et al., 2004). LFPs were low-pass filtered (cut-off frequency 400 Hz) and resampled at 2000 Hz using a FIR filter with optimal filter order. Noise due to the 60 Hz power-line frequency, as well as 30 Hz optical imaging interference and harmonic frequencies was removed using an adaptive filtering algorithm based on the Chronux toolbox functions (Bokil et al., 2010). Noisy contacts were identified using the power spectral density during spontaneous and evoked activity, signal to noise ratio (SNR; ratio of the power during evoked and spontaneous activity), and the correlation coefficient between adjacent channels along the probe. The signal from 1 to 2 non-adjacent electrode contacts was replaced with a signal interpolated over the neighboring channels using 2D splines. In practice, we had to replace noisy data from no more than 2 contacts in each experiment.

### 2.7. Seizure analysis

Seizures were identified and their number was estimated by visually inspecting the time courses of the recorded LFP and MUA. In addition to identifying seizures, we also estimated the seizure onset and termination times relative to the first pulse of the corresponding 10s-long stimulation period. Using the electrophysiological signals, the seizure onset time and duration were estimated for each of the seizures for each animal. Then, for each of the trials, including those with normal evoked responses, seizures, and refractory periods, we estimated the power of HFO ripples (80-200 Hz) and fast ripples (250-600 Hz). We estimated these power measures for three periods of each trial: the 5s before the onset of the stimulation period, during the stimulation period, and the 5s after the stimulation period.

### 2.8. Statistical evaluation

First, we tested if each of the samples represented a Gaussian distribution and whether the two samples had equal variances. In case either of these hypotheses was rejected, we followed with non-parametric testing; otherwise, we applied parametric testing. We used the Levene’s test to examine whether the variances of the compared groups were equal. In the cases where equal variances were verified, a Student’s t-test or post-hoc Tukey’s test was performed to test differences in the means of two or more groups, respectively. If there was evidence to support non-equal variances, a non-parametric Wilcoxon-Mann-Whitney’s test or Tamhane’s test were applied to evaluate statistical differences between two or more groups, respectively. A paired t-test was applied to groups of paired data variables. Results are presented as a mean ± SEM. Differences with p < 0.05 were considered statistically significant (IBM SPSS Statistics, 2016).

## 3. RESULTS

To examine whether peripheral electrical stimulation of the rat’s digit or forepaw can induce epileptic activity, we recorded LFP using a linear probe with 32 contacts inserted to the D3 representation in S1FL. The D3 representation was delineated based on optical imaging before inserting the probe and verified against the rat brain atlas.

### 3.1. Characterization of seizures

Figure 2A-C displays the LFP alongside the spectrogram of a seizure induced by somatosensory stimulation of the digit. As seen in Figure 2B, the onset (marked by a red arrow) of this seizure was associated with high-frequency negative or positive deflections (spikes) of the mean extracellular field potential. These events were later replaced by less frequent deflections with larger amplitudes extending above or below the baseline. These large-amplitude spikes appeared just before the seizure ended (Figure 2B).

To characterize the differences between normal response, seizure and refractory periods, Figure 3A presents an LFP time course that includes the three types of responses induced by digit D3 stimulation. A typical LFP pattern recorded during a normal evoked response is presented in Figure 3B. The neuronal responses were confined within the 10s period of stimulation. LFP amplitudes in response to electrical pulses were higher than the spontaneous LFP amplitudes at baseline, before stimulation began (Figure 2D). Figure 3C presents a time-course typical of the sensory stimulation induced seizure. Seizures were typically characterized by large amplitude spikes in the LFP and spike-waves throughout the duration of the epileptic activity (Figure 3C and 2D). Less frequently, rhythmic waves were observed. The seizures typically extended over several seconds beyond the stimulation period and were followed by refractory periods (10 refractory periods out of 17 seizures evoked by digit stimulation). During the refractory periods, a pattern of low-amplitude and slow oscillations was observed (Figure 3D and 2D; this time course is the average of 21 electrode contacts that spanned the cortical depth through all six layers). Less frequently, seizures were followed by a normal response (7 normal responses out of 17 seizures observed with digit stimulation).

The LFP responses induced by forepaw stimulation were similar to those induced by digit stimulation. Seizures induced by forepaw stimulation were similarly followed by a refractory period (40 refractory periods out of 113 seizures observed with forepaw stimulation) or a normal response (14 normal responses out of 113 seizures). However, in contrast to the events following digit stimulation, seizures induced by forepaw stimulation were followed most frequently by another seizure (55 cases out of 113 seizures). Refractory periods (n=40) were preceded by relatively long seizures (50.4±2.3s). Normal responses (n=14) were preceded by seizures of shorter duration (27.2±4.1s; p < 0.001, Tamhane’s test for comparing 3 classes, here comparing long and shorter seizures). On average, the first seizure in a pair of sequential seizures was short (17.8±1.2s; n=55; p < 0.001 between long and short seizures, Tamhane’s test).

### 3.2. Seizure susceptibility depends on anesthesia conditions and stimulation types

To determine the factors that influence seizure generation and spread, we applied different methods of anesthesia, recording, and stimulation applied to digit D3, digit D5, and forepaw. To evaluate the dependence of the seizure generation on the anesthesia regime, six experiments were performed using urethane, an anesthetic that has modest effects on both the inhibitory and excitatory systems and does not affect the noradrenergic system (Hara and Harris, 2002). As illustrated in Figure 4A-C, we did not evoke any seizure with D3 or D5 stimulation in any of the rats under urethane anesthesia.

**Figure 4.**
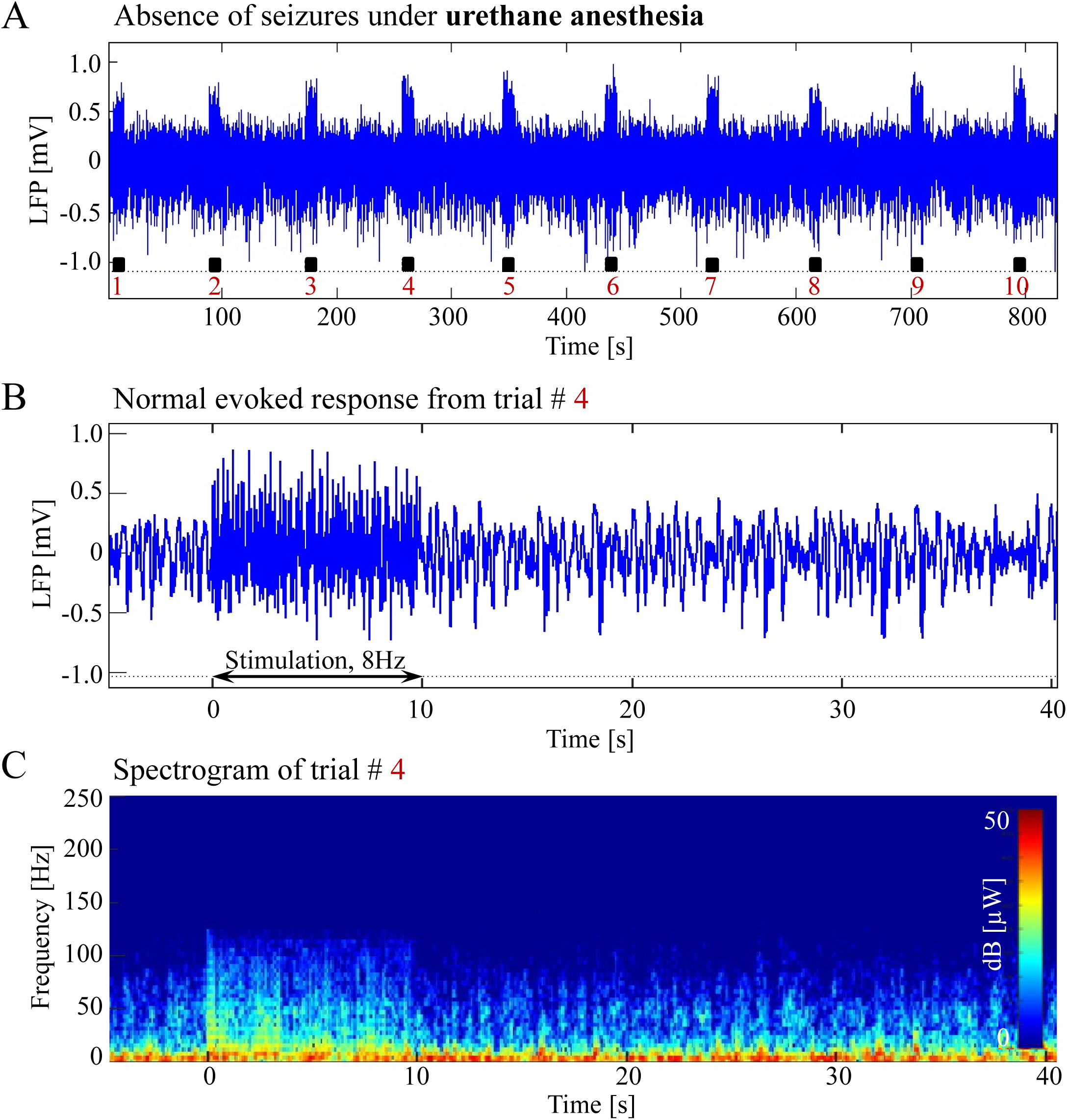
Absence of seizures under urethane anaesthesia. **A.** Ten trials of 10s-long D3 stimulation evoked no seizures following any of the stimulation periods. **B.** Normal LFP evoked responses to digit stimulation from the fourth trial. **C.** Spectrogram computed for the same trial, showing regular responses under urethane anaesthesia.

Neuronal responses in the somatosensory cortex constitute a spatiotemporal interplay between local excitation and surrounding inhibition (Cavanaugh et al., 2002; Derdikman et al., 2003). Derdikman et al. (2003) demonstrated triphasic responses consisting of an early depolarization, a late hyperpolarization, and a subsequent rebound depolarization. The region showing net hyperpolarization is on average more distant from the center of the responding area than the region showing net depolarization. To test whether seizures were more prone to occur right at the representation of the stimulated digit than in a neighboring region, we alternated stimulations of D3 and D5. With either stimulation of D3 or D5 (or forepaw, below), we recorded neurophysiological responses in the D3 cortical representation, based on the responses to D3 stimulation measured by optical imaging. On average, a larger number of seizures were detected following D3 stimulation than D5 stimulation; however, this was not statistically significant (p=0.27, paired t-test).

A balance between local excitation and surround inhibition is required to control the spread of local excitation over long distances (Derdikman et al., 2003). This raised the hypothesis that the seizures we demonstrated with dexmedetomidine and buprenorphine were elicited because our potent stimulation of a digit activated only a small area of cortex (single digit representation) which was insufficient to elicit a comparable surround inhibition. To test this hypothesis, in six rats we applied forepaw stimulation instead of the digit stimulation. The rationale was that forepaw stimulation activates large portions of area S1FL, and therefore it likely elicits more lateral inhibition than single digit stimulation (Remple et al., 2003; Waters et al., 1995). The LFP characteristics of seizures evoked by forepaw stimulation under dexmedetomidine and buprenorphine, including seizure onset and duration, were similar to those evoked by digit stimulation (Figure 5A-D; see details below). However, the percentage of rats that showed seizures during forepaw stimulation (60%) was higher than the corresponding percentage during digit stimulation (28%). In addition, the number of seizures during forepaw stimulation (18.83 ± 8.49 per animal) was higher than the corresponding number of seizures elicited by digit stimulation even though the means were not statistically significant (1.70 ± 0.70; Wilcoxon–Mann–Whitney test, Table 1).

**Figure 5.**
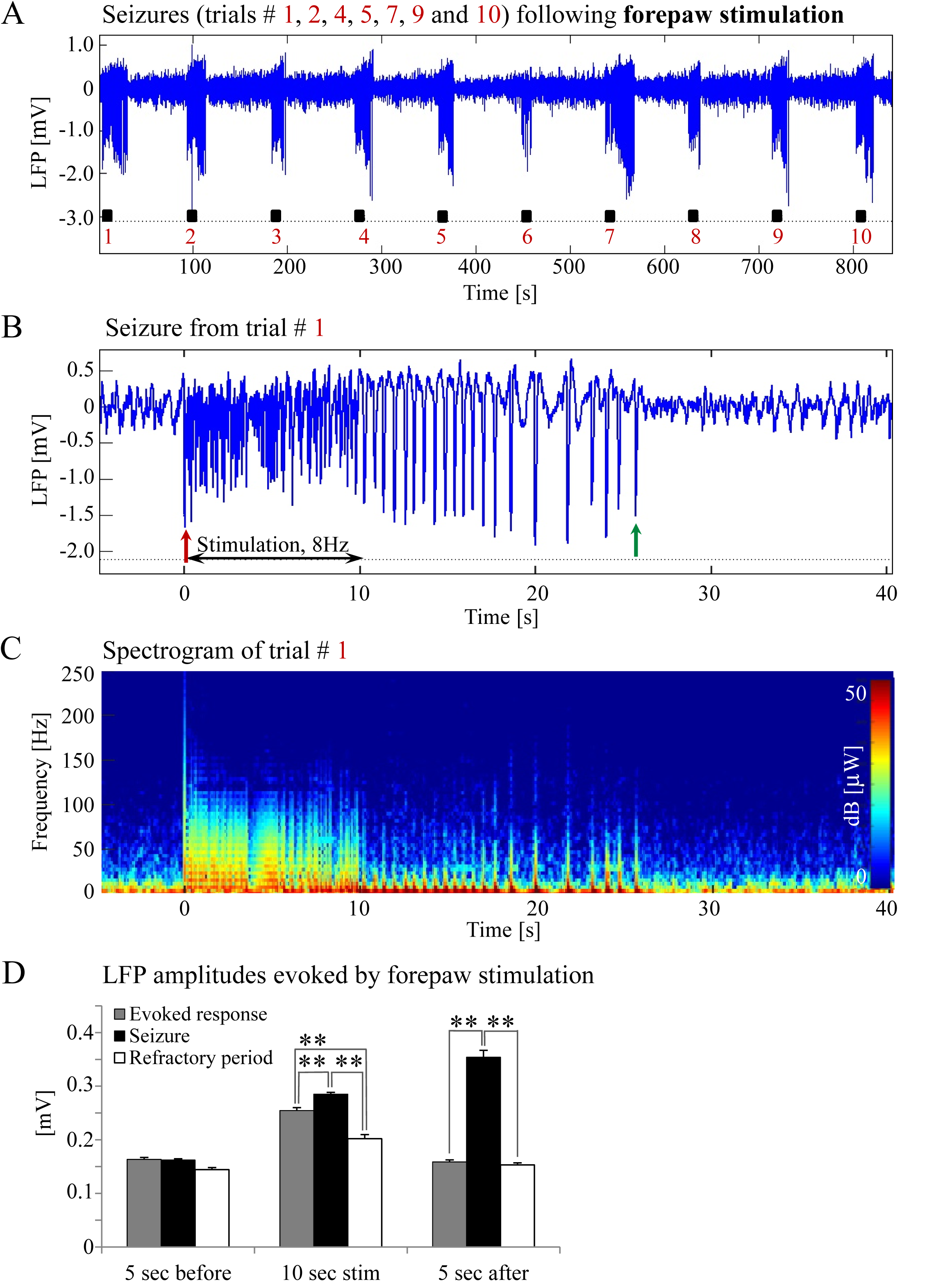
Seizures recorded with a linear probe following forepaw stimulation. **A.** LFP trace of ten trials, each with 10s-long stimulation. **B.** LFP (mean averaged over 20 electrode contacts spanning the cortical depth) recorded in one trial that showed a seizure pattern. The red and green arrows point to the onset and termination, respectively, of the seizure induced by forepaw stimulation. **C.** Spectrogram computed for the same seizure. **D.** The mean amplitudes of LFP recordings during baseline, stimulation and following stimulation for normal responses, seizures and refractory periods. The mean amplitudes of LFP represents the mean of the absolute mean extra-cellular potentials, obtained by averaging over probe contacts (** p<0.001; Tamhane’s test).

These results show that seizure vulnerability depends on the anesthesia regime and the type of peripheral stimulation. Sedation with dexmedetomidine and buprenorphine was associated with an increased probability of seizure induction relative to urethane anesthesia. Stimulating a larger cortical region by applying forepaw stimulation induced a larger number of seizures, relative to the number of seizures induced by digit stimulation.

### 3.3. Are the seizures caused by damage to cortex? Seizures recorded with surface ECoG arrays

It is well known that in humans and animals, mechanical damage to cortical brain areas can induce epileptic activity (Curia et al., 2016; Kharatishvili and Pitkanen, 2010). To examine whether damage caused by inserting the recording probe into the cortex had an effect on seizure generation, we performed additional experiments using surface ECoG arrays. These arrays were positioned on top of the dura matter; therefore, they did not cause any damage to cortex. Six rats under dexmedetomidine and buprenorphine sedation underwent forepaw stimulation. In five of six rats (84%) we observed seizures during and following forepaw stimulation (Figure 6A-D). The number of observed seizures per rat with ECoG recordings (24.16 ± 5.83) was higher than the number observed with intra-cortical probe (18.83 ± 8.49) and forepaw stimulation; however, this was not statistically significant (p=0.6, Student’s t-test). We concluded that the cortical damage caused by our intra-cortical probes was not a determining factor of seizure generation, since seizure susceptibility to forepaw stimulation was comparable across recordings with linear probes and ECoG arrays.

**Figure 6.**
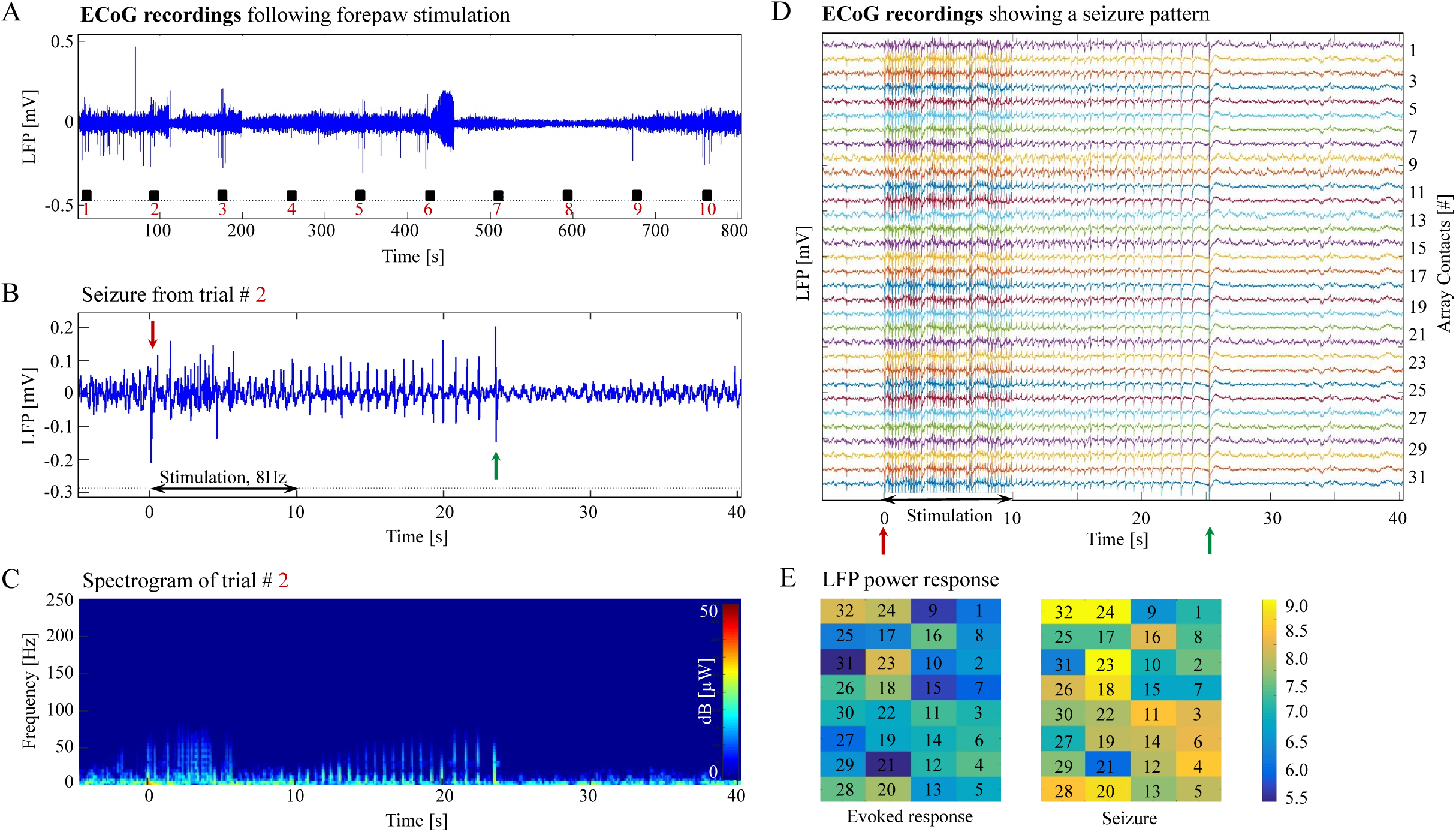
ECoG recordings detect epileptic activity induced by forepaw stimulation. **A.** A time-course averaged over all ECoG contacts: ten trials, each with 10s-long stimulation. **B.** ECoG recordings obtained in the second trial showing seizure activity recorded from the cortical surface. The red and green arrows indicate the onset and termination, respectively, of a seizure. **C.** Spectrogram computed for the same seizure**. D.** ECoG responses obtained from 32 surface contacts during one trial. **E.** Ratio of the power of the broad-band neurophysiological response during stimulation relative to the power during spontaneous activity. The left and right panels present this power ratio without (left) and with (right) seizure, respectively. The ratio is shown in an array form, presenting the power ratio obtained from each of the ECoG contacts in A. The channel numbers make it possible to relate the time-course in A to the pattern of the ECoG array in B.

### 3.4. Do the seizures propagate beyond the somatosensory representation of the forepaw?

Seizures often begin in an acute epileptic focus and spread tangentially to cortex, propagating to other brain regions (Li et al., 2018). As shown by Li et al. (2018) seizures induced by kainic acid injection into the hippocampus propagate to entorhinal and piriform cortex. To estimate the spatial extent of the seizures, we first analyzed the neuronal responses recorded by the ECoG array. The ratio between the LFP power during stimulation and baseline periods was calculated for each ECoG contact in order to visualize the normal response to forepaw stimulation (Figure 6E, left-side panel shows a normal response). We then repeated the same analysis for trials with induced seizures (Figure 6E, right-side panel shows seizure). ECoG array recordings showed that the normalized power of neuronal activity during a typical seizure was higher than that during the normal response in all contacts of the ECoG array (Figure 6D and E). Therefore, the data recorded by ECoG confirmed that the amplitudes of the ECoG signal are higher during seizures than during normal responses but could not determine differences in the spatial extent of these responses.

To test whether the spatial extent of the seizures is larger than that of the normal responses to digit and forepaw stimulation, we compared the spatial extent of the hemodynamic responses elicited during normal responses to forepaw stimulation to those elicited by seizures. Figures 7A and B present the spatial CBV responses evoked by stimulation of the forepaw. Note that the spatial extent of the active region immediately following stimulation seems slightly larger when a seizure takes place (Figure 7B, panel 2) than when the response is normal (Figure 7A, panel 2). The spatial extent of the active region 10s following the cessation of the stimulus is clearly larger when a seizure takes place (Figure 8B, panel 3) than when the response is normal (Figure 7A, panel 3). Two time-courses on the right display the corresponding temporal responses from the two brain regions within the activated brain area, which are marked on Figure 7C. At any time-point following the onset of the stimulus, the amplitude of the CBV response during the seizure (Figure 7B, red and blue time courses) were higher than the corresponding amplitudes during the normal evoked response (Figure 7A, corresponding time courses). The bar plots in Figure 7D present the spatial extent of the responses averaged over six animals, during normal responses and seizures, respectively. For each of the stimulation periods, the spatial extent of the cerebrovascular hemodynamic changes during seizures (Figure 7B, D) was larger than during normal evoked responses (Figure 7A and D). Note that the two latter periods, namely 10–15s and 15–20s relative to the onset of the stimulus, are part of the post stimulation period in which the seizures persist with no sensory stimulation n. Figure 8A, B and D shows similar differences between normal responses and seizures observed with a digit stimulation. We observed that during seizures the CBV increase extended ∼3 mm beyond the normal response, reaching the primary motor area (Figures 7B and 8B). Interestingly, not only the average spatial extent of the seizures was larger than that of normal responses obtained during the same period following stimulation, even the spatial extent during seizures *post stimulation* was larger than that observed in normal responses during stimulation (Figures 7D and 8D; p < 0.001, paired t-test). All animals displayed spatial spread of activity that surrounded the region of the D3 representation (for D3 stimulation) or of S1FL (for forepaw stimulation). Thus, the seizures we induced propagated beyond the spatial extent of the normal responses to forepaw stimulation to adjacent brain areas S1HL and part of M1, that were within the field of view of our optical imaging.

**Figure 7.**
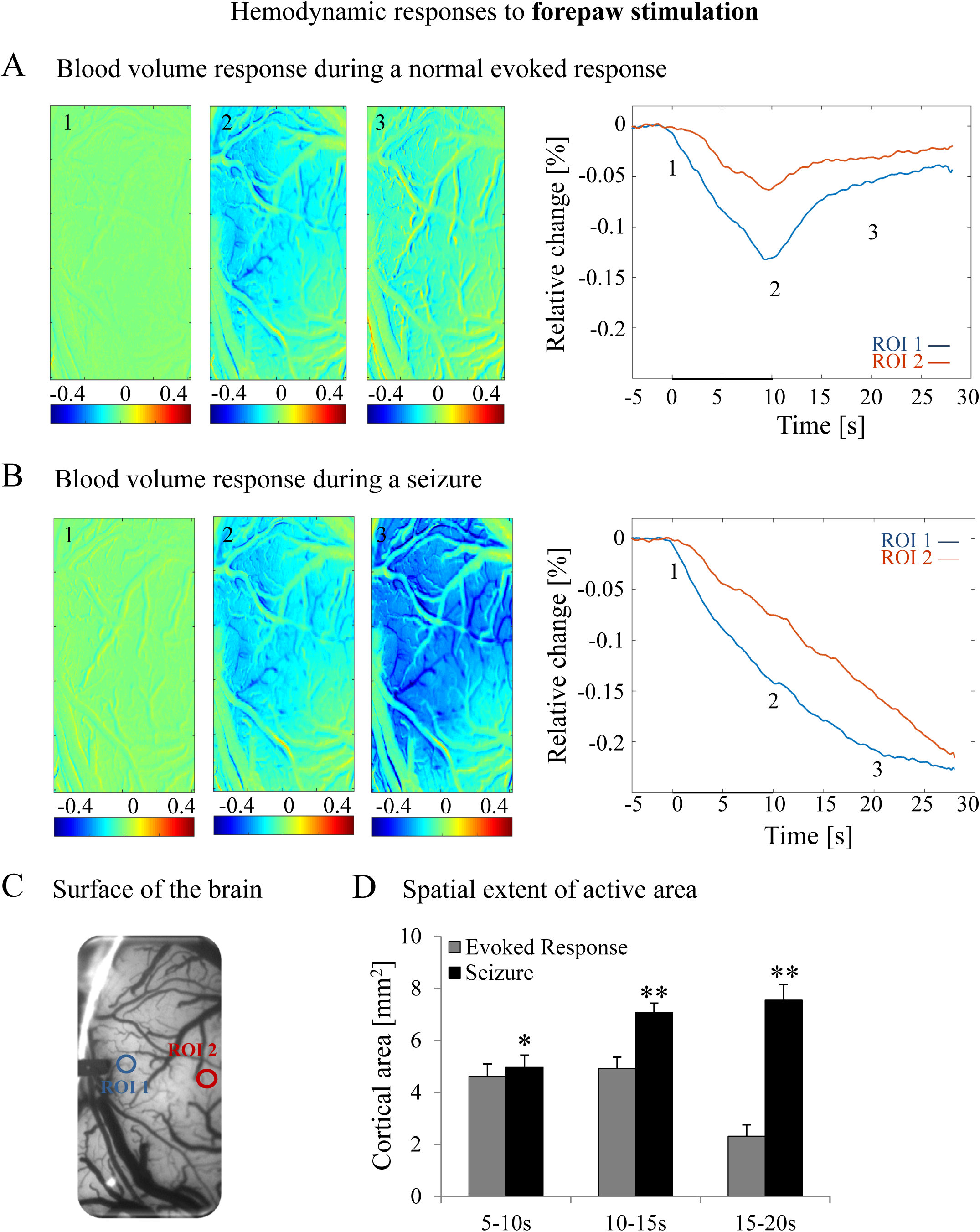
Cerebral blood volume response evoked by forepaw stimulation. **A.** Cerebral blood volume response evoked by stimulation of the contralateral forepaw. To the left, the spatial responses before (the 1s before stimulus onset), during (9-10s following stimulus onset), and after (19-20s following the stimulus onset) the 10s-long forepaw stimulation period. The reference for obtaining these responses was imaged between 3 and 1 seconds before stimulus onset. Note that negative response indicated in indexed blue color represent increase in absorption of light with wavelength 530 nm, i.e. increase in blood volume. To the right are two time-courses presenting the corresponding temporal response from two regions (blue and red ROIs in C) within the activated area. The stimulation period between 0 and 10 seconds is marked by a dark bar. **B.** Maps of the blood volume changes during seizure from before, during, and after the 10s-long forepaw stimulation period (exact time periods are as in A). To the right are two time-courses presenting the corresponding temporal response from two regions (blue and red ROIs in C) within the activated area. **C.** The imaged cortical surface with the two ROIs used for sampling the time-courses presented in A and B. **D.** A bar graph showing the spatial extent of the activated cortical region calculated for the epochs of 5-10s, 10-15s, and 15-20s relative to the onset of the stimulus during normal evoked responses (n=21) and seizures (n=21; * p<0.05, ** p<0.001; paired t-test between normal evoked responses and seizures). We calculated the spatial extent of the response trial by trial by applying a threshold (here, 3%) to the blood volume response in space. The same threshold was applied to all 3 periods of the normal evoked responses and seizures.

**Figure 8.**
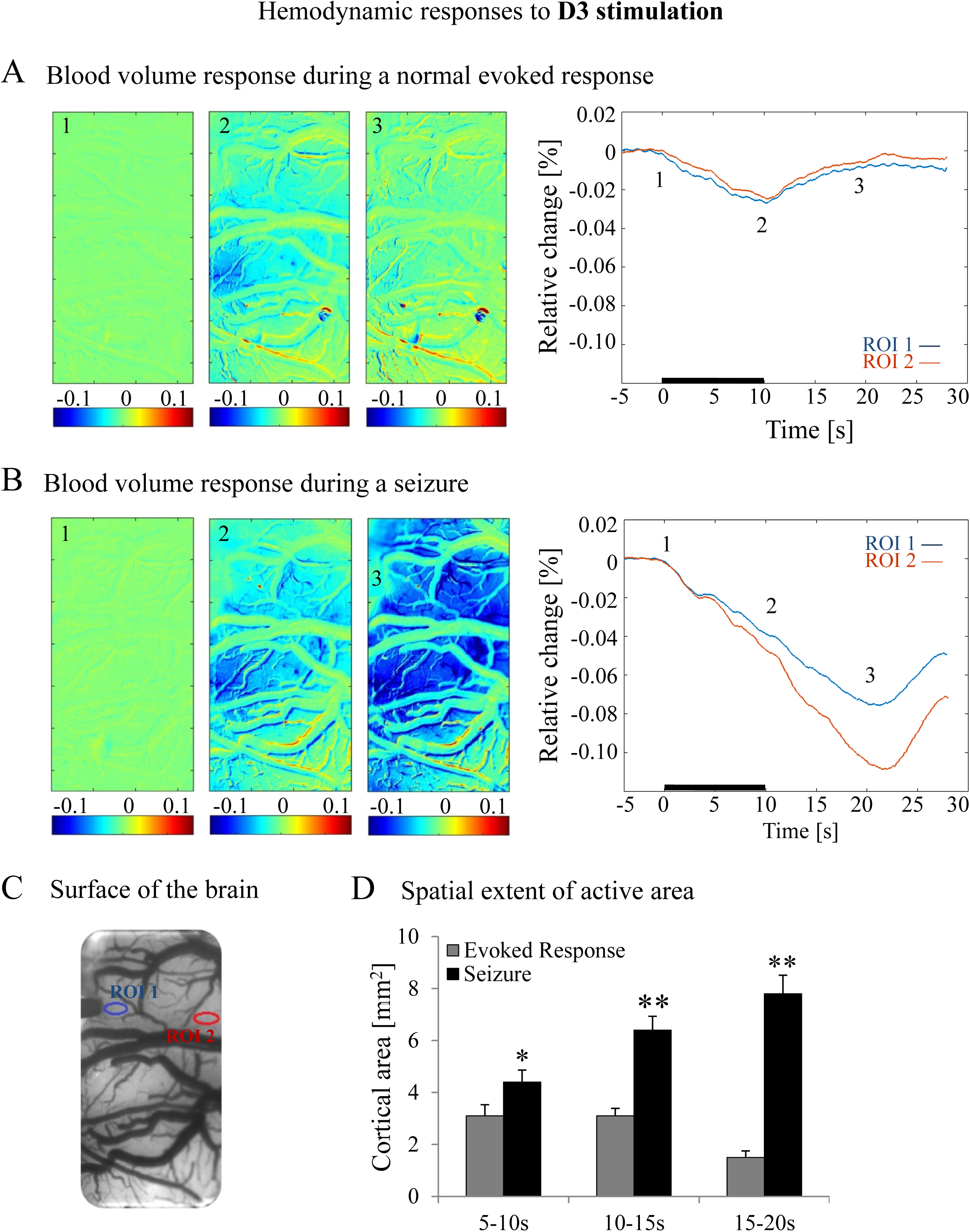
Cerebral blood volume response evoked by D3 stimulation. **A.** The format of presentation is identical to that used for Figure 7. In **D**, we calculated the spatial extent of the response trial by trial by applying a threshold (here, 2%) to the blood volume response in space (* p<0.05, ** p<0.001; paired t-test between normal evoked responses and seizures). The same threshold was applied to all 3 periods of the normal evoked responses and seizures.

### 3.5. Latency of the seizure onset and seizure duration

The characteristics of epileptic activity were evaluated for each group of rats. First, the seizure onset time and termination were marked by visual inspection (Figure 2). By examining the LFP amplitudes and the corresponding spectrograms, the seizure onset time was calculated relative to the first stimulation pulse. Analysis of the seizure onset time revealed no statistical differences between digit and forepaw stimulations and between recordings made with linear probe and ECoG array recordings (Table 1).

All seizures induced in our rats were brief, lasting less than one minute. Using linear probe recordings, the durations of seizures evoked by digit or forepaw stimulations were comparable. However, the average duration of seizures recorded with the ECoG array was shorter than that obtained with linear probe recordings (Table 1).

### 3.6. High-frequency oscillations

In the rat neocortex, HFOs can be observed in seizure-associated activity in the visual and sensorimotor cortex of animals (Jefferys et al., 2012; Murthy and Fetz, 1996). We therefore investigated whether HFOs coincided with the seizures in our animal model. Under dexmedetomidine and buprenorphine sedation, we observed ripples and fast ripples that preceded and coincided with all episodes of seizures induced by peripheral somatosensory stimulation (Figure 9A-B). The power of fast ripples during the 5s prior to stimulation associated with seizures was higher relative to the power observed prior to normal responses and refractory periods (Figure 9D). The power of fast ripples following the end of stimulations associated with seizures was higher than the corresponding power associated with normal evoked responses and refractory periods (Figure 9D).

**Figure 9.**
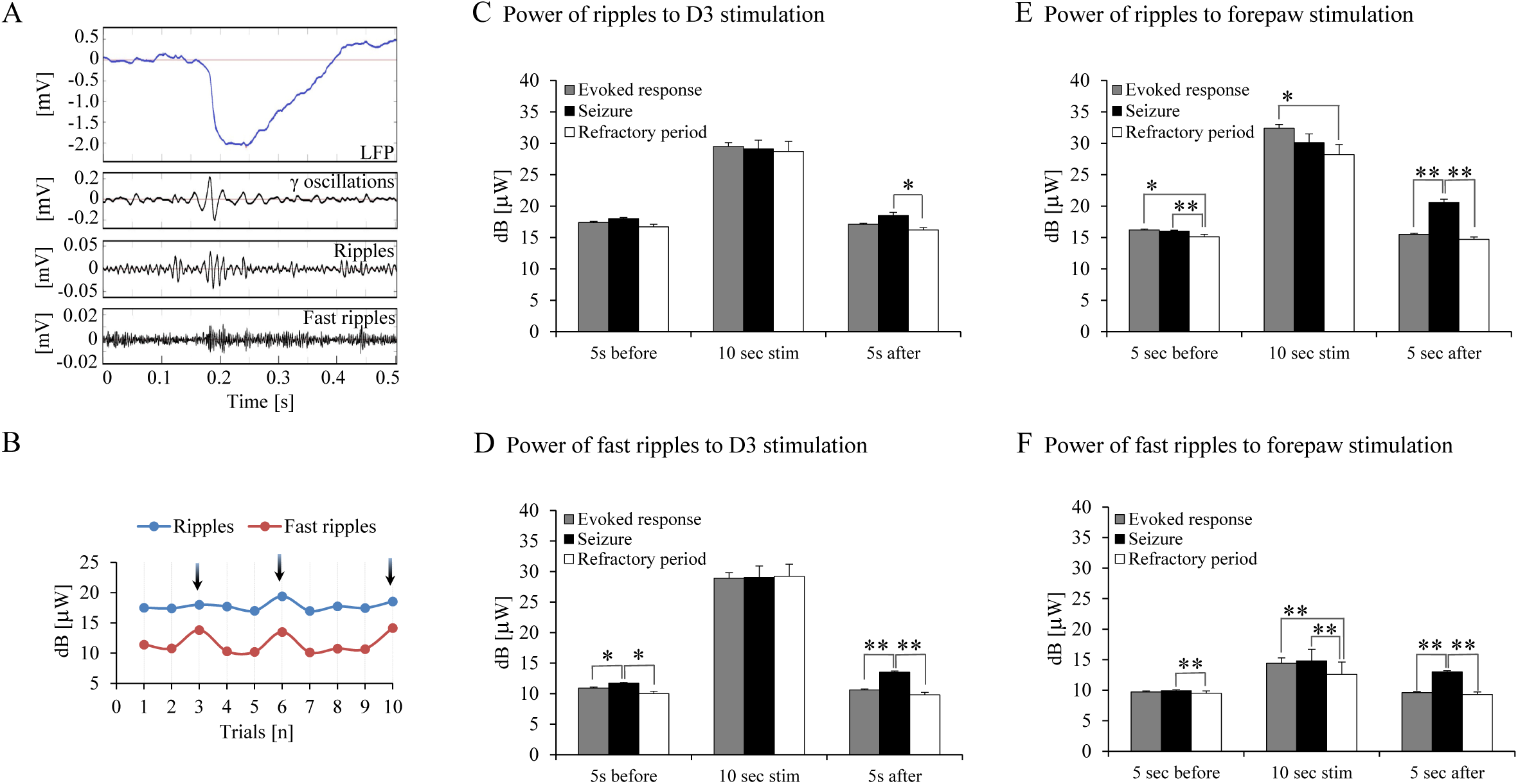
HFOs in response to D3 and forepaw peripheral somatosensory stimulation. **A.** LFP recorded from area S1FL showing an after-discharge with simultaneously occurring ripples (80-200 Hz) and fast ripples (250-600 Hz) following stimulation of D3. High frequency oscillations coincided with all episodes of seizure induced by digit or forepaw stimulation. **B.** A representative example of the power of ripples and fast ripples corresponding to the 5s period after the cessation of stimulation. The black arrows indicate the trials with seizures. Similar variations in the power of HFOs were observed in all animals following digit or forepaw stimulation. **C.** A bar graph showing the average power of the ripples to D3 stimulation 5s before, during, and 5s after the stimulations associated with normal evoked responses (n=129 trials), seizures (n=21 trials), and refractory periods (n=20 trials). Results are shown as a mean ± SEM. Note the statistically significant difference between seizures and refractory periods (* p<0.05; Tamhane’s test) following the offset of the stimulation. **D.** Average power of the fast ripples 5s before, during, and 5s after the stimulation associated with normal evoked responses (n=129 trials), seizures (n=21 trials), and refractory periods (n=20 trials). Statistically significant differences were observed between normal evoked responses and seizures before and after the stimulation period, and between seizures and refractory periods before and after the stimulation period (* p<0.05, ** p<0.001; Tamhane’s test). **E.** A bar graph showing the average power of the ripples to forepaw stimulation 5s before, during, and 5s after the stimulations associated with normal evoked responses (n=50 trials), seizures (n=110 trials), and refractory periods (n=50 trials). Results are shown as a mean ± SEM. Note the statistically significant differences between evoked responses and refractory periods, evoked responses and seizures, and seizures and refractory periods (* p<0.05; ** p<0.001; Tamhane’s test). **F.** Average power of the fast ripples 5s before, during, and 5s after the stimulation associated with normal evoked responses (n=50 trials), seizures (n=110 trials), and refractory periods (n=50 trials). Statistically significant differences were observed between evoked responses and refractory periods, evoked responses and seizures, and seizures and refractory periods (** p<0.001; Tamhane’s test).

With forepaw stimulation, the power of ripples during seizures was higher than the corresponding power during refractory periods before and after the stimulation period (Figure 9E). The power of ripples following seizures was higher than the power observed following normal evoked responses after the stimulation period (Figure 9E). The power of fast ripples before seizures was higher than the power observed before responses during the refractory period (Figure 9F). The power of fast ripples during the stimulation periods associated with evoked responses and seizures was higher than the power observed during stimulation during the refractory period (Figure 9F). The power of fast ripples following seizures was higher than the power observed following normal evoked responses and following responses during refractory periods (Figure 9F).

Lastly, for normal evoked responses, seizures, and refractory periods, the power of ripples and fast ripples during stimulation was higher than the power observed during the 5s periods before and after the stimulation with digits (p < 0.001; Tamhane’s test; Figure 9C-D, bars in the middle) and forepaws (p < 0.001; Tamhane’s test; Figure 9E-F).

## 4. DISCUSSION

Based on the experiments presented here, we have described five main findings. First, electrical stimulation administered peripherally to the rat digits D3 or D5, as well as to the forepaw, produces seizures. Second, seizure susceptibility depends on anesthesia conditions: sedation with dexmedetomidine increases the probability of seizure induction in comparison to anesthesia with urethane. Third, seizure susceptibility also depends on the spatial extent of the stimulated region, i.e. single digit or forepaw representation. Fourth, the evoked seizures propagate to adjacent brain cortical regions, beyond the spatial representation of the stimulated digit or forepaw. Fifth, high-frequency oscillations precede and coincide with all seizures evoked by peripheral stimulation. Importantly, the seizures we induced by peripheral electrical stimulation form a new animal model of somatosensory-induced reflex seizures. This model is free of electrical artifacts and does not involve damage to the brain caused by the insertion of a stimulating electrode or application of chemo-convulsant.

### 4.1. Characteristics of the induced seizures

Our findings indicate that electrical somatosensory stimulation of naïve rat digits or forepaw forms an animal model of reflex seizures. In epileptic patients, TIS can be evoked by a cutaneous-mucosal sensorial stimulus and is limited to specific skin regions (Sala-Padro et al., 2015; Striano et al., 2012). Focal areas that are sensitive to these stimuli include the foot, finger, arm, shoulder, chest, face, and head (Deonna, 1998). In majority of patients affected by TIS, the sensory stimulations that induce seizures are repetitive and last 5-20 seconds (Hsieh et al., 2011; Kanemoto et al., 2001; Sala-Padro et al., 2015). To trigger the epileptic activity in our animals, we followed a similar approach to somatosensory stimulation. The electrical stimuli we delivered to the digit or forepaw were repetitive and lasted 10 seconds. Another feature of our seizure model similar to TIS in humans is that TIS are brief and frequently followed by a refractory period (Deonna, 1998; Hsieh et al., 2011; Sala-Padro et al., 2015; Striano et al., 2012). All our animals displayed similar, brief electrographic seizures that were followed by a refractory period, normal response, or another seizure.

The neocortex is one of the brain structures involved in several types of acquired epilepsy, including TIS (Chauvette et al., 2016; Timofeev et al., 2014). Cortical seizures are characterized by spike– and polyspike–wave electrographic after-discharges, as well as fast oscillations that are followed by the formation of larger amplitude spike– and polyspike– wave field potential complexes (Timofeev et al., 2014). The spike-and-wave after-discharges were also observed in cat S1 during direct stimulation of the thalamic ventrobasal complex (Dietzel et al., 1982; Mares et al., 1984). A similar pattern of epileptiform electrographic activity is observed in our animal model of somatosensory-induced reflex seizures (e.g., Figure 2B). The repetitive nature of stimulation we applied is reminiscent of kindling procedures. Kindling is commonly induced by the application of repetitive, focal, sub-convulsive electrical stimulation directly to specific brain structures (such as the amygdala or hippocampus), leading to after-discharges and electrographic seizures (Barnes and Pinel, 2001; Goddard et al., 1969; Gorter et al., 2016; Morimoto et al., 1987). After-discharges can be triggered by electrical stimulation administered directly to different brain regions, including cortical area S1 (Dietzel et al., 1982; Kalamangalam et al., 2014; Lesser et al., 2008; Mares et al., 1984). In contrast to our proposed model, the process of kindling is laborious and time consuming, requiring precise electrode implantation and electrical stimulation that can extend over days (Raol and Brooks-Kayal, 2012). Importantly, kindling is induced by direct stimulation of the brain’s tissue, whereas the seizures we present here are induced by peripheral sensory stimulation.

In an epileptic brain, cortical seizures elicit a focal, large-amplitude increase in utilization of glucose and oxygen, cerebral blood vessel dilatation, and an increase in blood flow and volume (DeSalvo et al., 2010; Engel et al., 1982; Lauritzen and Gold, 2003; Patel et al., 2013). Traditional models of seizure propagation involve the assumption of an onset zone and a progressive outward spread of activation (Ma et al., 2013). Neuroimaging techniques, such as intrinsic optical imaging, indirectly measure neuronal activity based on changes in CBV and oxygenation. Intrinsic optical imaging under illumination of an isosbestic wavelength such as 530 nm is directly proportional to increases and decreases in CBV (Ma et al., 2013; Zhao et al., 2009). As shown in previous studies, intrinsic optical signal can be used to identify ictal initiation and early propagation (Bahar et al., 2006; Zhao et al., 2009). In our animal model, all epileptic events induced by digit or forepaw stimulation affected the CBV. All seizures began focally and propagated horizontally through the cortex. The entire S1FL region and parts of surrounding cortical areas displayed a steady increase in CBV (Figures 7B and 8B). The spatial extent of the CBV response to the seizures was larger than the S1FL representation of the stimulated digit or forepaw. In addition, the spatial extent of the CBV responses to seizures increased with time, confirming that seizures are prone to cortical spread. The epileptic responses we elicited show characteristics similar to those demonstrated by Haglund and Hochman (2007) using optical imaging of intrinsic signals, where seizures were induced by focal application of bicuculine to the cortical surface. In addition, similar hemodynamic responses were observed during seizures evoked by focal administration of penicillin directly onto cortex (Chen et al., 2000), bicuculine (Suh et al., 2005) and 4-aminopyridine (Harris et al., 2014; Ma et al., 2013). Lastly, in humans affected by neocortical epilepsy, epileptiform events increase CBV and the signal spreads widely throughout the cortex (Shariff et al., 2006), mirroring our animal seizure data.

### 4.2. Characteristics of the anaesthetic regime and stimulation type

In the present study, seizures were evoked under a combination of a sedative – dexmedetomidine - and the morphine-related analgesic buprenorphine. The strong opioid analgesic effect of buprenorphine should exclude the pain-related pathway of seizure induction. Accordingly, the animal physiological parameters, including heart rate and oxygen saturation were not affected by the peripheral electrical stimulation and we did not observe any motor response such as contraction of muscles that could indicate pain during stimulation. This type of sedation possesses neuroprotective characteristics such as optimized cerebral oxygen supply and decreased intracranial pressure and CBV during surgery (Bekker and Sturaitis, 2005; Gerlach and Dasta, 2007). It has been reported that under continuous intravascular systemic perfusion of dexmedetomidine for more than two hours, forelimb stimulation elicits seizure-like responses observed in LFP, accompanied by changes in cerebral blood flow (Fukuda et al., 2013). Indeed, it was suggested that the use of dexmedetomidine *per se* modulates central adrenergic function by activating presynaptic α2-adrenergic receptors (Gertler et al., 2001), which in turn increases sensitivity to seizure expression (Oishi and Suenaga, 1982). In contrast, Whittington et al. (2002) demonstrated that dexmedetomidine suppresses the induction, generalization, and severity of rat seizures induced by intravenous administration of cocaine. Moreover, it was reported that medetomidine sedation does not increase seizure vulnerability, nor does it affect LFP and BOLD responses during epileptic activity (Airaksinen et al., 2012). Consistent with the findings by Whittington et al. (2002) and Airaksinen et al. (2012), we did not evoke any epileptic activities with short forelimb stimuli protocols (1s stimulation of 8 Hz with 0.6-0.8 mA) under the same anesthetic condition (Sotero et al., 2015). Therefore, based on data of Whittington et al. (2002), Airaksinen et al. (2012), and our results (Sotero et al., 2015), dexmedetomidine sedation is not sufficient for inducing epileptic activity with short peripheral stimulation. To induce seizures, we had to use long (10 s) and potent (2 mA) peripheral electrical stimuli (Figure 2).

To elucidate whether the seizures depend on the dexmedetomidine sedation regime, we applied the same stimulation paradigm under a different anesthesia regime, using urethane. Urethane is an anesthetic that produces modest effects on both inhibitory and excitatory systems and has anticonvulsive properties (White et al., 1954). The changes exerted on multiple neurotransmitter-gated ion channels are much smaller than those seen with anesthetics more selective for one neurotransmitter system, such as ketamine. Urethane, therefore, is suitable for maintaining anesthesia during electrophysiological recording (Hara and Harris, 2002). Under urethane anesthesia, we did not evoke any seizure. Similarly, Heltovics et al. (1995) applied 3-aminopyridine topically to the surface of cortex and showed that urethane completely abolishes epileptic activity induced by this chemo-convulsant and prevents the development of recurrent seizures. In addition, although kainic acid typically causes convulsions, urethane anesthesia limits the effects of systemic kainic acid application to only electrographic seizures with no convulsions (Saito et al., 2006).

The seizures observed with ECoG were shorter and had a slightly late onset time as compared with intra-cortical probe and forepaw stimulation. This difference can be explained by the linear probes’ capacity to detect activity manifested by increases in high frequencies – such as spiking activity – which is better than that of ECOG’s. Linear probes are inserted into cortex and the surface area of their contacts is smaller than those of ECoG arrays. Their contacts are in close proximity to the neurons. In contrast, ECoG arrays were positioned on top the dura matter. Their sensitivity to high-frequencies is substantially reduced, because the surface area of their contacts is larger than that of the probes’, and because cortex conducts high-frequencies to shorter distances than the distances over which it conducts low-frequencies. This explanation also holds for why high-frequency activity during baseline can be better observed with intra-cortical probes but only to a lesser extent with ECoG arrays. Therefore, while we cannot rule out that the damage caused by inserting the probes to cortex exacerbated the seizures, it is unlikely that this damage was a determining factor in the generation of seizures.

Seizures induced by forepaw stimulation were more frequent than those induced by digit stimulation under similar stimulation parameters. The somatosensory cortical representation of the forepaw is larger than that of individual digits (Coq and Xerri, 2000; Remple et al., 2003; Waters et al., 1995). Therefore, our findings suggest that the likelihood of generating seizures increases with the spatial extent of the stimulated cortical tissue.

### 4.3. High frequency oscillations as biomarkers of epileptogenic neural networks

Pathological HFOs can be recorded in the LFP of epileptic patients (Jacobs et al., 2010; Malinowska et al., 2015; Perucca et al., 2013) as well as in animal epilepsy models (Bragin et al., 2004; Levesque et al., 2011). Earlier studies have shown that seizure onset zones can be identified by HFOs (Modur et al., 2011; Ochi et al., 2007; Salami et al., 2014). Similarly, our animal model shows HFOs within the epileptic onset zone and can be used for studying the role of HFOs in seizure generation.

In the rat neocortex, spontaneous, somatosensory-evoked and pathological HFOs have been observed (Jones and Barth, 1999; Staba, 2012). As described by Jones and Barth (1999), the sensory-evoked neocortical HFOs occur in conjunction with the biphasic positive-negative somatosensory-evoked potential. Similarly, our findings show that the power of ripples and fast ripples increases during normal somatosensory-evoked responses (Figure 9). It was also shown that the power of HFOs dramatically increases at the onset as well as during seizures (Grenier et al., 2003; Jiruska et al., 2010). Likewise, we show that HFOs recorded in our animals precede episodes of seizures and HFO power increases during seizures induced by peripheral somatosensory stimulation (Figure 9). This indicates that the corresponding neural network is more excitable and therefore, more prone to evoke seizures.

### 4.4. An artifact-free animal model of reflex seizures

Electrical stimulation models were the first prototypes introduced for studying seizures, status epilepticus, and neuronal excitability (Loscher, 2011; Reddy and Kuruba, 2013). External electrical stimulation protocols are less harmful to subjects and offer better control of seizures compared to chemical induction methods (Kandratavicius et al., 2014; Loscher, 2011). However, the majority of external electrical stimulation methods of seizure induction generate seizures in large parts of the brain, rather than focal seizures. Electrical stimulation administered directly in the brain causes damage to the tissue. In addition, electrical stimulation methods cause large artifacts which interfere with electrical recordings and disable investigation of the spatiotemporal mechanisms of seizure induction (Osawa et al., 2013).

We introduce an animal model of somatosensory-induced reflex seizures that represents brief and artifact-free, non-invasively induced seizures, with no application of damage, current or convulsant directly to the brain. As shown by our optical imaging measurements, the initial seizures propagate from their onset zones to other cortical regions. Consequently, our model can be used for studying the mechanisms of seizure propagation. The model of seizures we propose is simple to use; it neither involves animal mortality, nor it require sophisticated equipment. Epileptic activity in this animal model is evoked by non-invasive, peripheral somatosensory electrical stimulation. This allows a more natural way for inducing focal seizures compared to the invasive methods used currently, which involve direct application of drugs to the brain. Our proposed animal model elicits focal seizures that propagate following the seizure onset. Our proposed model can be used to shed light on the characteristics of the onset, propagation, maintenance, and termination of seizures in a lamina-specific manner.

## 5. CONCLUSION

Our study reveals that the seizures we induced by peripheral electrical stimulation of rat digits or forepaws form a new animal model of somatosensory-induced reflex seizures for studying seizure generation and propagation and therapeutic approaches of human reflex epilepsy. This model is free of electrical artifacts and does not involve damage to the brain caused by the insertion of a stimulating electrode or application of chemoconvulsants. Based on our data, forepaw electrical somatosensory stimulation induces seizures more reliably and at a higher rate than digit stimulation. Therefore, we recommend forepaw stimulation when using this animal model of seizures.

## Acknowledgements

This study was supported by the Canadian Institute of Health Research (Grant MOP-102599) and the Natural Sciences and Engineering Research Council of Canada (RGPIN 375457-09 and RGPIN 2015-05103) awarded to A.S. We thank Roland Pilgram for the pipeline for analyzing electrophysiological data, and Jean Gotman, Massimo Avoli, Victor Mocanu and Pascal Kropf for their helpful technical advice and discussions. We thank Laura Diamond and Katie Hu for scientific English editing.

## Notes

#### Summary of Updates

Objective: We introduce a novel animal model of somatosensory stimulation-induced reflex seizures which generates focal seizures without causing damage to the brain. Methods: Specifically, we electrically stimulated digits or forepaws of adult rats sedated with dexmedetomidine while imaging cerebral blood volume and recording neurophysiological activity in cortical area S1FL. For the recordings, we either inserted a linear probe into the D3 digit representation or we performed surface electrocorticography (ECoG) recordings. Results: Peripheral stimulation of a digit or the forepaw elicited seizures that were followed by a refractory period with decreased neuronal activity, or another seizure or normal response. LFP amplitudes in response to electrical pulses during the seizures (0.28 +- 0.03 mV) were higher than during normal evoked responses (0.25+- 0.05 mV) and refractory periods (0.2 +- 0.08 mV). Seizures generated during the stimulation period showed prolonged after-discharges that were sustained for 20.9 +- 1.9 s following the cessation of the stimulus. High-frequency oscillations were observed prior to and during the seizures, with amplitudes higher than those associated with normal evoked responses. The seizures were initially focal. Optical imaging of the cerebral blood volume response showed that they propagated from the onset zone to adjacent cortical areas, beyond the S1FL representation of the stimulated digit or forepaw. The spatial extent during seizures was on average 1.74 times larger during the stimulation and 4.1 times following its cessation relative to normal evoked responses. Seizures were recorded not only by probes inserted into cortex but also with ECoG arrays (24.1 +- 5.8 seizures per rat) placed over the dura matter, indicating that the seizures were not induced by damage caused by inserting the probes to the cortex. Stimulation of the forepaw elicited more seizures (18.8 +- 8.5 seizures per rat) than stimulation of a digit (1.7 +- 0.7). Unlike rats sedated with dexmedetomidine, rats anesthetized with urethane showed no seizures, indicating that the seizures may depend on the use of the mild sedative dexmedetomidine. Significance: Our proposed animal model generates seizures induced by electrical sensory stimulation free of artifacts and brain damage. It can be used for studying the mechanisms underlying the generation and propagation of reflex seizures and for evaluating antiepileptic drugs.

